# Liver Enzymes and Blood Lactate Profile of Patients Diagnosed with Typhoid Fever in Abuja, Nigeria

**DOI:** 10.1101/2022.11.19.517198

**Authors:** Oluchukwu Ogechukwu Anunobi, Rita Ewela Ojo

## Abstract

**Background:** Typhoid fever resistance is on the increase and this frustrates efforts at treatment. Persistence of multiple drug-resistant (MDR) typhoid fever leads to higher mortality rate since treatment is evasive.

**Objectives:** This study intended to identify if liver enzymes and lactate levels are related to aggravating MDR in patients diagnosed with typhoid fever.

**Methods:** 50 subjects were recruited, 45 were positive for Widal test and further subjected to stool culture examination for the presence of *S*. Typhi. All subjects blood were analysed for lactate, alanine aminotransferase (ALT), aspartate aminotransferase (AST), and alkaline phosphatase (ALP)

**Results:** Growth of *S*. Typhi was observed in only eight subjects out of the 45 Widal positive test subjects. The eight positive *S*. Typhi isolates showed resistance to the first line drugs, macrolides and 3^rd^ generation cephalosporins but showed susceptibility to fluoroquinolones. ALP, AST, ALT and blood lactate were elevated in all multi-drug resistant patients. Highest levels of liver enzymes were observed in subjects resistant to the greatest number of antibiotics. However, there was no significant correlation of increase in blood lactate level and ALP when compared with increase in resistance based on number of antibiotics (R^2^=0.2858, R^2^=0.4675).

**Conclusion:** Lactate and liver enzyme levels increase markedly during typhoid fever infection, however in MDR scenario, especially in resistance to different classes of antibiotics, the metabolic toll on the liver increases and is reflected in elevation of AST and ALT.

## INTRODUCTION

Typhoid fever is an endemic infection caused by *Salmonella* serotype Typhi. It is acquired through oral-faecal transmission. It is often diagnosed through blood, urine, stool and bone marrow culture.^(1)^ In Nigeria, typhoid fever remains a major disease burden because of factors such as increased urbanization without adequate supplies of portable water, inadequate facilities for processing human waste, overburdened health-care delivery systems, and overuse of antibiotics that contribute to the development and spread of resistant strains of *S*. Typhi. However, the true incidence of typhoid fever is difficult to evaluate in Nigeria because of the lack of a proper coordinated epidemiological surveillance system. Nevertheless, information on typhoid fever prevalence has been reported by several researchers in some states in Nigeria ranging from 0.071% in Oyo to 47.1% in Osun.^(2)^ Despite the advent of successful typhoid fever vaccines, majority of the population are ignorant of it, while most cannot afford it.

To cope with the several antibiotic use as therapeutics against *S*. Typhi, the bacterium has developed defenses against some of these antimicrobials employed. The degree of resistance against the antibiotics could be against one or more, of which resistance to at least three first-line antibiotics is referred to as multi-drug resistant.^(3)^ Despite the development of new improved antibiotics that are effective against *S*. Typhi resistance to most of the front-line antibiotics is growing more prevalent globally. The isolation of multi-drug resistant strains and the more recent class called the extensive-drug resistant strains from clinical samples is on the rise.^(4)^ It is estimated that there is about 14.3 million typhoid fever infections and more than 135,000 deaths worldwide annually.^(5)^ There are several mechanisms that could be responsible for *S*. Typhi resistance to antibiotics, and this includes enzymatic inhibition, alteration of bacterial membranes, rapid drug efflux, reduced drug influx, introduction of new drug resistant target and alteration of target sites. Genetic mutation or horizontal gene transfer could also cause antibiotic resistance. This alteration of the genome of the organism could be an adaptation strategy and thus fosters the evolution of these strains into more resilient species.^(6, 7)^ Studies have shown that drug resistance in *S*. Typhi is mostly due to a genetic modification in the organism, either as a chromosomal mutation or the acquisition of a plasmid or transposon. ^(8, 9, 10)^

Elevated lactate levels have been observed among typhoid fever patients. However, the presence of elevated lactate levels is seen to manifest in most sepsis-related diseases and its presence in typhoid fever is not unexpected, especially because it is related to hepatomegaly and some mildly deranged liver function.^(11)^ In this study we investigated the levels of lactate and some liver enzyme in different types of antimicrobial resistance observed in typhoid fever outpatients.

## METHODS

### Study area and population

Study population included male and female outpatients aged 7 - 75 years reporting to Karu General Hospital Abuja, laboratory from May-June, 2019. Sample size for this study was 50 samples.

### Ethical consideration

This study and procedures followed were in accordance with the ethical standards of the of National Health Research Ethics on human experimentation and the Helsinki Declaration of 1975, as revised in the year 2013. the ethical approval (BHU/REC/19/H/0001) was obtained from Research and Ethics Committee Bingham University, Karu, Nigeria. Written informed consent to publication was obtained from patients before inclusion into the study.

### Experimental Design

Patients directed to the laboratory by the consulting doctor for Widal tests were given written informed consent forms and questionnaires after explaining the research to them. A semistructured questionnaire was employed in obtaining information from participants in this study about their age, sex, duration of illness, pre-treatment information, and their disease recurrence rate. Trained phlebotomist collected blood samples from patients who complied and part of the blood sample was used for Widal test. Only patient samples positive for Widal test were considered eligible and further analysed for lactate and select liver enzymes. Eligible patients were given stool sample bottles to bring freshly-passes morning stool the next day. Routine typhoid fever test (Widal test) was carried out and healthy individuals not suspected to have typhoid fever were recruited as control.

### Sample Collection

Using standard phlebotomy techniques, 5 ml of venous blood was collected from patients. 3ml transferred to Ethylene Diamine Tetra-Acetic Acid (EDTA) containers and gently mixed, while 2ml was transferred to lithium heparin bottle. Freshly passed stool was also collected into sterile sample containers and labelled accordingly.

### Widal test

Blood collected in EDTA bottle was centrifuged and sera obtained was used in Widal test to identify significant serum agglutins titer against typhoid/paratyphoid antigens (according to manufacturer’s protocol).

### Blood Lactate Analysis

An amount of 0.5 ml of whole blood taken from the lithium heparin bottle was used to analyse lactate levels using rapid Accutrend Plus diagnostic meter and BM Lactate test strips (Roche Diagnostics). Accutrend Plus diagnostic meter employs colorimetric lactate oxidase/mediator reaction and has a record of high reproducibility when compared with laboratory diagnosis.^(12)^

### Liver enzymes analysis

1.5ml of blood sample in lithium heparin bottle was centrifuged at 3000rpm for 3 minutes and then 1 ml of sera was pipetted into sample cups and used to analyse for ALT, AST and ALP levels using an automated biochemical analyser (Selectra pro S 6003-500) applying the principle of colorimetry.

### Stool sample collection and culture

Freshly passed early morning stool was collected in sterile wide mouth sample containers. Stool specimen was inoculated into Selenite F broth in McCartney bottles, and kept for 18 hours at 37°C to encourage selective enrichment of *Salmonella spp*. After 18 hours, the inoculum was streaked directly onto Salmonella-Shigella Agar and incubated at 37°C for 24 hours. The resulting colonies were observed for colorless colonies with black centre and sub-cultured to obtain pure cultures.

### Identification of isolates

Isolates were identified based on their cultural characteristics, Gram stain reaction, cell morphology, slide agglutination and biochemical tests like triple sugar iron, urease test and lack of oxidase.^(13)^

### Antibiotics Susceptibility Testing

The antibiotics susceptibility testing was done by Kirby-Bauer disk diffusion method according to NCCLS guidelines using 30 mcg of amoxicillin, 25 mcg of Cotrimoxazole, 5 mcg of Levofloxacin, 5 mcg of Ofloxacin, 30 mcg of Tetracycline, 30 mcg of Netillin, 10 mcg of Gentamicin and 30 mcg of Ceftriaxone. A colony of the test organism was picked using a sterile wire loop and emulsified in a sterile bottle using normal saline. A sterile non-toxic cotton swab on a wooden applicator was dipped into the inoculum and the soaked swab rotated to drain excess liquid. The entire Muller Hinton agar surface of the plate was streaked with the swab containing the inoculum and allowed to dry for 5-10 minutes with the lid in place. Using sterile forceps, the antimicrobial disks were then placed on the agar. The plate was incubated at 37°C for 18 to 48 hours. Using a ruler on the underside of the plate, the radius of each inhibition zone was measured in millimeters and the value doubled to get the diameter. The zones of inhibition were read and results interpreted using standardized thresholds for defining susceptibility. ^(14)^

### Statistical analysis

Antimicrobial susceptibility frequency data was presented in percentages. Lactate and liver enzymes analysis data was presented as Mean (SD) and compared with theoretical/hypothetical values using the Student t-test. Pearson’s correlation was employed to identify any correspondence between antimicrobial resistance and lactate and liver enzymes levels. P<0.05 was considered statistically significant. All data were statistically evaluated using Graphpad Prism 7.

## RESULTS

Out of the 50 patients tested, 32% were male while 68% were female. Subjects within the age range of 20-40 years had the highest cases of Widal positives (58%). Out of the 45 Widal positives, prevalence of *S*. Typhi in patient’s stool was 8 isolates (18%) and 37 isolates (82%) were identified as negative for growth of *Salmonella* ser. Typhi in their stool. The 37 negatives had growth of other bacteria like *Proteus mirabilis, E*.*Coli, Citrobacter freundii* etc in their stool. The five patients that served as control did not show any observable growth in their stool culture.

Data obtained from the patient’s questionnaires on disease duration, disease frequency, pre-treatment and dosage of drugs are shown in Table 1.

**Table 1:** Frequency of demographics/data from *S*. Typhi positive subject’s questionnaires

| Data | Subjects (n=8) |
| --- | --- |
| Disease frequency $\geq 3$ times | 8 (100%) |
| Disease frequency $< 3$ times | 0 (0%) |
| Disease duration $\geq 3$ weeks | 7 (87.5%) |
| Disease duration $< 3$ week | 1 (12.5%) |
| Self-treatment with local remedy | 2 (25%) |
| Self-treatment with ethno-medicinal drugs | 6 (75%) |
| Adherence to drug regimen | 4 (50%) |
| Inconsistency to drug regimen | 4 (50%) |

The antibiotics resistance rates in Table 2 shows that 75% of the stool culture positive for *S*. Typhi were resistant to Tetracycline, 62.5% resistant to Ceftriaxone, 100% resistant to amoxicillin, 87.5% resistant to Cotrimoxazole, 87.5% resistant to Gentamicin, 67.5% to Netillin, 12.5% resistant to Levofloxacin and 0% resistant to Ofloxacin.

**Table 2:** Frequency of Antimicrobial resistance patterns of *S*. Typhi positive patients (n=8)

| S/N | Antimicrobial agent | Disk<br>potency<br>(mcg) | Sensitive | Intermediate | Resistant |
| --- | --- | --- | --- | --- | --- |
| 1. | Tetracycline | 30 | 1 (12.5%) | 1 (12.5%) | 6 (75%) |
| 2. | Ofloxacin | 5 | 7 (87.5%) | 1 (12.5%) | 0 (0%) |
| 3. | Ceftriaxone | 30 | 2(25%) | 1 (12.5%) | 5 (62.5%) |
| 4. | Levofloxacin | 5 | 7(87.5%) | 0 (0%) | 1 (12.5%) |
| 5. | Amoxicillin | 30 | 0 (0%) | 0 (0%) | 8 (100%) |
| 6 | Cotrimoxazole | 25 | 0 (0%) | 1 (25.5%) | 7 (87.5%) |
| 7. | Gentamicin | 10 | 0 (0%) | 1 (12.5%) | 7 (87.5%) |
| 8. | Netillin | 30 | 3 (37.5%) | 0 (0%) | 5 (62.5%) |

The lactate values, AST, ALP, ALT and antibiotics resistance type in *S*. Typhi positive subjects are depicted in Table 3. The mean value for lactate was 10.5(2.24) mmol/L, 65.15(4.96)U/L for ALT, 55.08(6.78)U/L for AST and 158.10(8.32)U/l for ALP.

**Table 3:** Lactate, ALT, AST, ALP and antibiotics resistance in the *S*. Typhi positive isolates

| S/N | Age | Sex | Lactate value<br>(1.8–2.2mmol/L) | ALT<br>(7–56U/L) | AST<br>(10–40U/L) | ALP<br>(20–120IU/L) | Antibiotics Resistance | No. of Resistant Drugs |
| --- | --- | --- | --- | --- | --- | --- | --- | --- |
| 0001 | 35 | F | 6.0 | 62.4 | 49.2 | 149.0 | Multi-drug | 4 |
| 0003 | 31 | F | 10.8 | 63.7 | 55.9 | 161.2 | Multi-drug | 5 |
| 0004 | 37 | F | 12.0 | 69.3 | 59.8 | 163.0 | Multi-drug | 6 |
| 0005 | 40 | M | 9.1 | 59.5 | 52.3 | 157.9 | Multi-drug | 5 |
| 0010 | 50 | F | 13.6 | 71.2 | 61.8 | 165.5 | Multi-drug | 6 |
| 0011 | 45 | M | 12.3 | 69.7 | 61.0 | 163.7 | Multi-drug | 6 |
| 0023 | 10 | F | 11.2 | 67.4 | 58.4 | 162.4 | Multi-drug | 5 |
| 0038 | 47 | F | 9.0 | 58.0 | 42.2 | 141.9 | Multi-drug | 2 |

## DISCUSSION

In most developing countries like Nigeria, typhoid fever is widely diagnosed with only Widal test and clinical features and this formed the basis of our inclusion criteria. Typhoid fever cases reported in this study were first diagnosed using Widal test before the stool culture analysis. The absence of *S*. Typhi growth in the stool of 82% of the patients after testing positive for Widal test could be due to cross reacting antibodies from serum of febrile patients other than typhoid fever, ^(15)^ and other infections sharing a common antigenic determinant with *S*. Typhi,^(16)^ due to low specificity of Widal tests.^(17)^ It is also possible that some were in the clinical relapse phase, which refers to the return of a disease or its symptoms after a period of improvement. This was stated in a report by Miller and Peuges ^(18)^ who observed that clinical relapse caused absence of growth of *S*. Typhi in the stool.

Resistance to the first line drugs (Amoxicillin, Tetracycline and Cotrimoxazole), macrolides (Gentamicin and Netillin) and third generation cephalosporins (Ceftriaxone) was observed in all eight *S*. Typhi positive isolates. Maximum resistance was observed in amoxicillin; 8 (100%) isolates were resistant, Cotrimoxazole; 7(87.5%) isolates were resistant, 1(12.5%) isolate was intermediately susceptible and with Gentamicin; 7(87.5%) isolates were resistant and 1(12.5%) isolate was intermediately susceptible. The fluoroquinolones (Ofloxacin and

Levofloxacin) were seen to be effective treatments with more susceptibility rates. Ofloxacin showed higher efficiency in 7(87.5%) isolates and intermediate susceptibility seen in 1(12.5%) isolate. 7(87.5%) isolates showed susceptibility to Levofloxacin with only 1(12.5%) isolate resistant to it. This is consistent with the report of Alam ^(19)^ who observed high susceptibility to Levofloxacin and Ofloxacin (Table 2). Similar trend has also been reported by Willey ^(20)^ where high resistance to first line drugs, macrolides and 3^rd^ generation cephalosporins and susceptibility to fluoroquinolones were observed. The multiple drug resistance observed in all *S*.Typhi positive isolates could be due to abuse of these drugs. Self-treatment by patients for typhoid fever when they have other febrile diseases such as non-typhoidal Salmonellosis, endocarditis and urinary tract infection in turn results to development of resistance. ^(21)^ Misdiagnosis of typhoid fever in its absence also leads to misuse of the antimicrobials. The abuse and misuse of these antimicrobials consistently makes the bacteria develop defense mechanisms, such that the bacterium becomes resistant to the intended drug effect.

Data obtained from *S*. Typhi positive patient’s questionnaire (Table 1) revealed that they have had typhoid fever for 3 weeks or more, and evidently supported by the presence of *S*. Typhi in the stool as *S*. Typh*i* is excreted in the stool 2-3 weeks post infection ^(22)^. All *S*. Typhi positive patients underwent self-medication using various antibiotics, which included Amoxicillin, Ceftriazone, and Tetracycline in a bid to treat the disease, of which 50% didn’t complete the self-treatment regimen. Those drugs may have been abused by constant ingestion irrespective of being prescribed or not and this could have led to the development of resistance. ^(23)^ The antibiotics resistant typhoid fever seen in 75% of the eight isolates could also be as a result of pre-treatment with ethno-medicinal therapy (Table 1). Fifty percent of *S*. Typhi positive subjects who claim to have always adhered to their drug regime but still were resistant could be as a result of contracting an already resistant strain of *S*. Typhi to the antibiotics in question. Multi-drug resistance to some antibiotics observed in 50% of patients that failed to adhere to their drug regime could be because of incomplete dosage of drugs which causes the destruction of some bacteria while the undestroyed ones go on to develop more resistance properties Elevated lactate levels we observed in *S*. Typhi positive patients could be due to typhoid fever as well as other factors (Table 4) since prevalence of elevated lactate levels was observed in all *S*.Typhi positive subjects with a mean concentration of 10.5(2.24) mmol/L way above the normal range of 1.8-2.2 mmol/L. The highest lactate levels were observed in the patient with highest number of drug resistance. There was a significant difference observed when mean lactate levels of *S*.Typhi positive subjects were compared with theoretical normal. It is not a surprise either that the lactate levels of some of the 37 patients who were negative for *S*.Typhi culture were also elevated (Table 6). Most of these patients may be suffering other ailments like malaria which is also known to elevate blood lactate levels. ^(24)^

**Table 4:** Comparison of mean Lactate, AST, ALT and ALP values with control

| Parameter | Actual mean | Control mean | P value | t, df |
| --- | --- | --- | --- | --- |
| **Lactate (mmol/L)** | 10.50(2.40) | 1.58(0.52) | P<0.0001 | t=10.54, df= 7 |
| **AST (UI/L)** | 55.08(6.78) | 33.30(0.67) | P<0.0001 | t=9.076 df=7 |
| **ALT (UI/L)** | 65.15(4.96) | 45.50(2.25) | P<0.0001 | t=11.2 df=7 |
| **ALP (UI/L)** | 158.10(8.32) | 84.25(8.69) | P<0.0001 | t=25.09 df=7 |

There may not be adequate reports on elevation of lactate levels in typhoid fever, however it was discovered and reported by Sameera ^(25)^ that there is a statistically significant increase in lactate dehydrogenase (an enzyme that catalyzes the conversion of pyruvate to lactate) in typhoid fever patients when compared with healthy controls (Suppl. 1). This elevated lactate dehydrogenase levels in typhoid fever cases corroborates the usefulness of lactate levels data in the determination of clinical and prognostic feature of typhoid fever.

Typhoid fever results in an increase in lactate levels when the cytotoxins from *S*. Typhi cause necrosis to lymphoid tissues and injury to the kupffer cells of the liver. When this occurs, *S*. Typhi release endotoxins that cause toxemia. ^(26)^ As the liver is clearly implicated in this process, it is not surprising to find liver enzymes elevated during and active typhoid infection.

Antibiotics resistance showed negative correlation with lactate levels (R^2^=0.2858) (Table 5). Although there is biochemical explanation for elevated lactate in typhoid fever, we didn’t observe a direct relationship between increasing antibiotics resistance and increasing lactate levels. It was observed in this study that lactate level did not increase accordingly with an increase in antibiotics resistance. The elevated lactate levels observed could just be due to expected dysfunction of the liver during active typhoid fever infection.

**Table 5:** Correlation Analysis between Lactate, Antibiotics Resistance, ALT, AST and ALP.

|  | **Resistance**<br>**Vs.**<br>**Lactate** | **Resistance**<br>**Vs.**<br>**ALT** | **Resistance**<br>**Vs.**<br>**AST** | **Resistance**<br>**Vs.**<br>**ALP** |
| --- | --- | --- | --- | --- |
| **Pearson's correlation coefficient** | 0.5346 | 0.9083 | 0.8036 | 0.5346 |
| **R<sup>2</sup>** | 0.2858 | 0.8249 | 0.6458 | 0.4675 |
| **P value** | 0.1722 | 0.0018 | 0.0163 | 0.0615 |
| **P value summary** | ns | ** | * | ns |

**Table 6:**
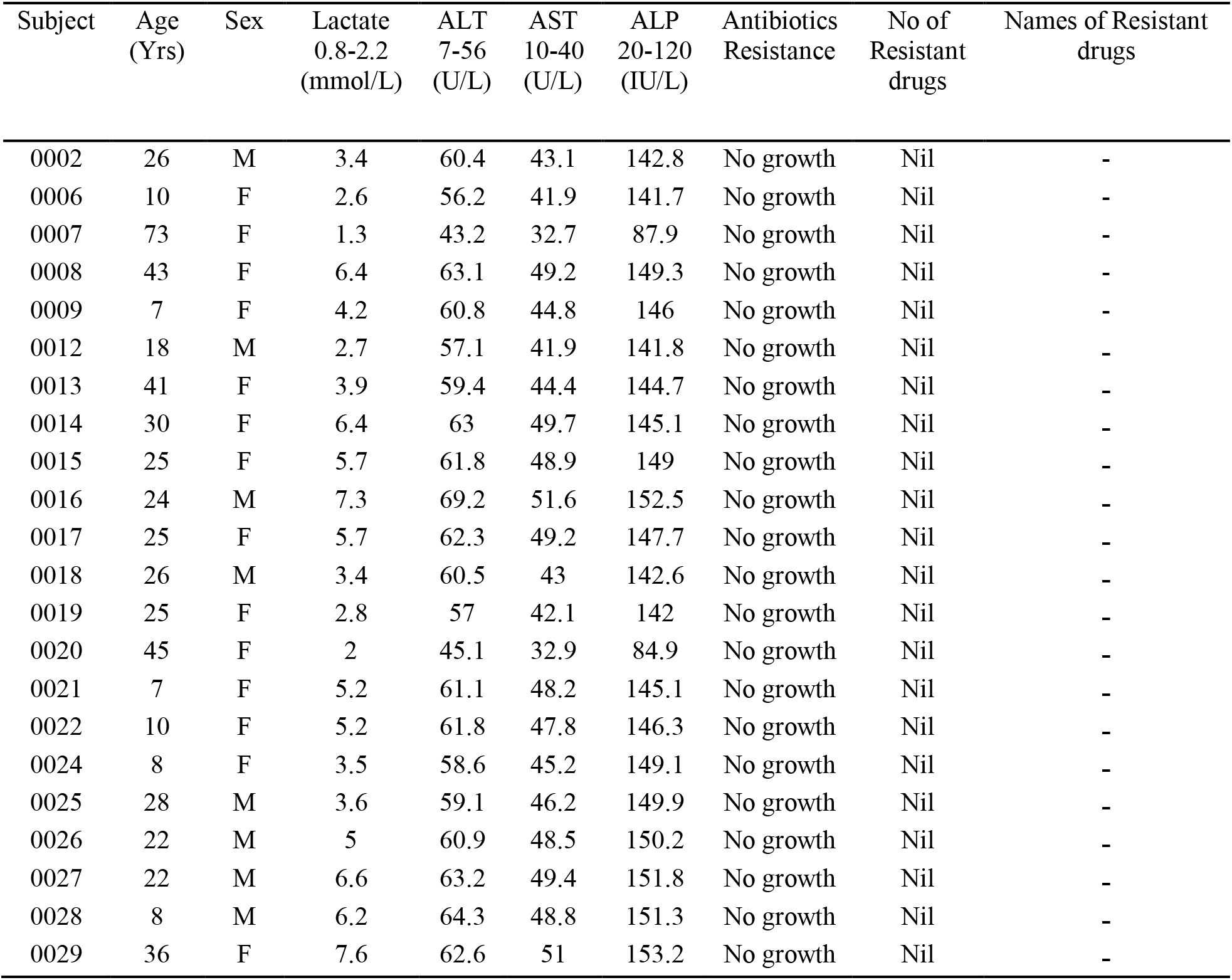
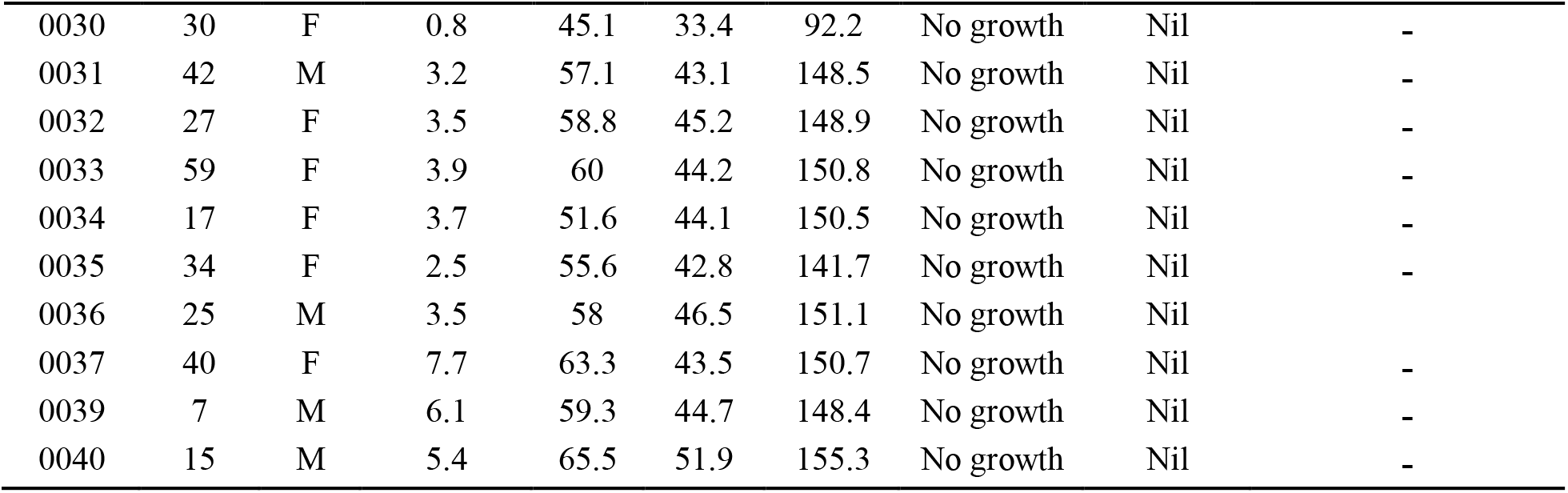
Data on all parameters analysed in all negative *S*. Typhi culture subjects

| Subject | Age<br>(Yrs) | Sex | Lactate<br>0.8-2.2<br>(mmol/L) | ALT<br>7-56<br>(U/L) | AST<br>10-40<br>(U/L) | ALP<br>20-120<br>(IU/L) | Antibiotics<br>Resistance | No of<br>Resistant<br>drugs | Names of Resistant<br>drugs |
| --- | --- | --- | --- | --- | --- | --- | --- | --- | --- |
| 0002 | 26 | M | 3.4 | 60.4 | 43.1 | 142.8 | No growth | Nil | - |
| 0006 | 10 | F | 2.6 | 56.2 | 41.9 | 141.7 | No growth | Nil | - |
| 0007 | 73 | F | 1.3 | 43.2 | 32.7 | 87.9 | No growth | Nil | - |
| 0008 | 43 | F | 6.4 | 63.1 | 49.2 | 149.3 | No growth | Nil | - |
| 0009 | 7 | F | 4.2 | 60.8 | 44.8 | 146 | No growth | Nil | - |
| 0012 | 18 | M | 2.7 | 57.1 | 41.9 | 141.8 | No growth | Nil | - |
| 0013 | 41 | F | 3.9 | 59.4 | 44.4 | 144.7 | No growth | Nil | - |
| 0014 | 30 | F | 6.4 | 63 | 49.7 | 145.1 | No growth | Nil | - |
| 0015 | 25 | F | 5.7 | 61.8 | 48.9 | 149 | No growth | Nil | - |
| 0016 | 24 | M | 7.3 | 69.2 | 51.6 | 152.5 | No growth | Nil | - |
| 0017 | 25 | F | 5.7 | 62.3 | 49.2 | 147.7 | No growth | Nil | - |
| 0018 | 26 | M | 3.4 | 60.5 | 43 | 142.6 | No growth | Nil | - |
| 0019 | 25 | F | 2.8 | 57 | 42.1 | 142 | No growth | Nil | - |
| 0020 | 45 | F | 2 | 45.1 | 32.9 | 84.9 | No growth | Nil | - |
| 0021 | 7 | F | 5.2 | 61.1 | 48.2 | 145.1 | No growth | Nil | - |
| 0022 | 10 | F | 5.2 | 61.8 | 47.8 | 146.3 | No growth | Nil | - |
| 0024 | 8 | F | 3.5 | 58.6 | 45.2 | 149.1 | No growth | Nil | - |
| 0025 | 28 | M | 3.6 | 59.1 | 46.2 | 149.9 | No growth | Nil | - |
| 0026 | 22 | M | 5 | 60.9 | 48.5 | 150.2 | No growth | Nil | - |
| 0027 | 22 | M | 6.6 | 63.2 | 49.4 | 151.8 | No growth | Nil | - |
| 0028 | 8 | M | 6.2 | 64.3 | 48.8 | 151.3 | No growth | Nil | - |
| 0029 | 36 | F | 7.6 | 62.6 | 51 | 153.2 | No growth | Nil | - |
| 0030 | 30 | F | 0.8 | 45.1 | 33.4 | 92.2 | No growth | Nil | - |
| 0031 | 42 | M | 3.2 | 57.1 | 43.1 | 148.5 | No growth | Nil | - |
| 0032 | 27 | F | 3.5 | 58.8 | 45.2 | 148.9 | No growth | Nil | - |
| 0033 | 59 | F | 3.9 | 60 | 44.2 | 150.8 | No growth | Nil | - |
| 0034 | 17 | F | 3.7 | 51.6 | 44.1 | 150.5 | No growth | Nil | - |
| 0035 | 34 | F | 2.5 | 55.6 | 42.8 | 141.7 | No growth | Nil | - |
| 0036 | 25 | M | 3.5 | 58 | 46.5 | 151.1 | No growth | Nil | - |
| 0037 | 40 | F | 7.7 | 63.3 | 43.5 | 150.7 | No growth | Nil | - |
| 0039 | 7 | M | 6.1 | 59.3 | 44.7 | 148.4 | No growth | Nil | - |
| 0040 | 15 | M | 5.4 | 65.5 | 51.9 | 155.3 | No growth | Nil | - |

Alanine amino transaminase (ALT), Aspartate amino transaminase (AST) and Alkaline phosphatase (ALP) are referred to as diagnostic liver enzymes because in elevated states, can be indicative of liver malfunction. Studies have associated elevated liver enzymes with active typhoid fever infection. ^(27, 28)^ As expected with most liver related dysfunctions, AST 55.08(6.78) U/L, ALT 65.15 (4.96) U/L and ALP 158.10(8.32) U/L, were elevated in subjects with active typhoid infection. There was a significant increase observed in the comparison of mean ALT (P<0.0001), AST (P<0.0001) and ALP (P<0.0001) levels of *S*. Typhi positive patients with their respective control values (Table 4). This increase in liver enzymes may be due to hepatomegaly caused by endotoxins produced from *S*. Typhi and direct invasion of the hepatocytes by *S*. Typhi. A large population of *S*. Typhi attach themselves to the kupffer cells such that kupffer cells which are phagocytic are unable to destroy the bacteria population. Invasion of *S*. Typhi in the liver causes injuries and release of liver enzymes in large quantities. Hepatomegaly is not a complication of typhoid fever but a feature of the disease. ^(29)^

Increase in antibiotics resistance had a positive correlation with the ALT (R^2^=0.8249) and AST (R^2^=0.6458) showing that higher concentrations of AST and ALT in typhoid fever may indicate the severity of the disease. Resistance to antibiotics therapy will result in an untreated condition of typhoid fever and in this active infectious condition, the liver is burdened. The higher the number of drugs the pathogen is resistant to, the longer the infection persist without treatment and more severe the infection.

Since no significant correlation was observed between antibiotics resistance and ALP (R^2^=0.4675) concentration although ALP concentrations were elevated in all S. Typhi positive subjects, it shows no dependence of any on the other (Table 5). Increased ALP concentration was not dependent on increase in antibiotics resistance.

## CONCLUSION

Further abuse and misuse of antibiotics for treatment of typhoid fever will continue to disarm our therapeutic arsenal of antimicrobials until total resistance is observed. Unfortunately, health officers who prescribe these antimicrobials using only Widal test positive result for diagnosis are contributing to this increased resistance and this is still practiced widely in Nigeria. Elevated blood lactate levels take a metabolic toll on the body and should be considered in every case of typhoid fever even though its increase does not depend on the level of resistance the *S*. Typhi manifests. AST and ALT levels on the other hand may be indication that a patient is a MDR *S*. Typhi carrier and prompt an antibiotics susceptibility test to guide prescription of drugs.

## Acknowledgements

We would like to thank Odiba Solomon Ph.D. and Olugbodi Janet Ph.D. for their assistance and guidance in this research.

---

**Figure 1:**
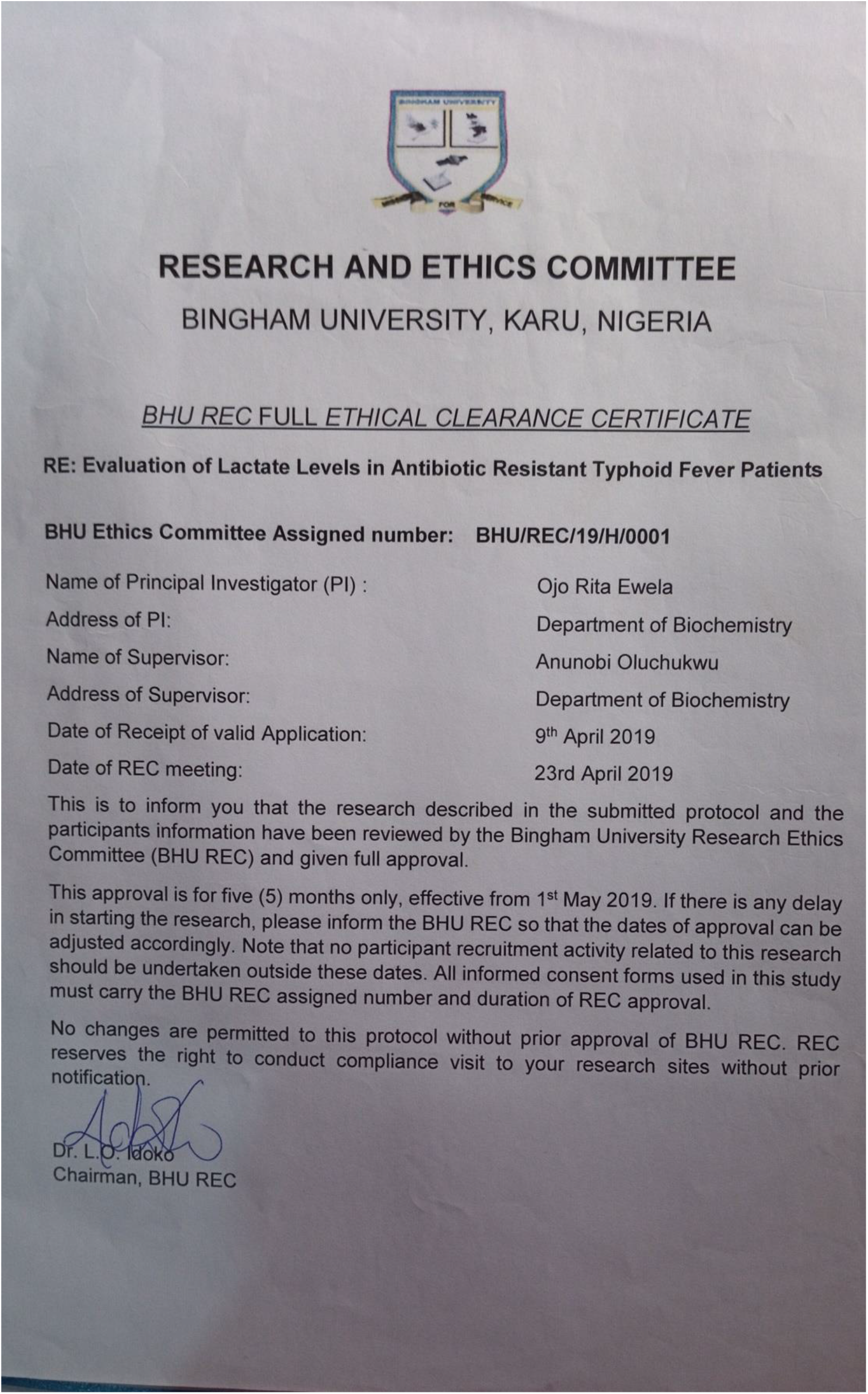
Ethical approval document.

