## Supplemental Table 1 for "Liver Enzymes and Blood Lactate Profile of Patients Diagnosed with Typhoid Fever in Abuja, Nigeria"

Supplemental table 1: Data on all parameters analysed in all normal subjects (control)

| Subject | Age (Yrs) | Sex | Lactate 0.8-2.2 (mmol/L) | ALT 7-56 (U/L) | AST 10-40 (U/L) | ALP 20-120 (IU/L) | Antibiotics Resistance | No of Resistant drugs |
|---|---|---|---|---|---|---|---|---|
| 0046 | 24 | F | 2 | 63.2 | 48.7 | 121.4 | No growth | Nil |
| 0047 | 32 | F | 1.8 | 48.6 | 34.2 | 72 | No growth | Nil |
| 0048 | 24 | M | 1.3 | 58.2 | 45.1 | 112.8 | No growth | Nil |
| 0049 | 35 | M | 1.3 | 53.8 | 45.5 | 101.8 | No growth | Nil |
| 0050 | 21 | M | 0.8 | 49.1 | 45.3 | 121.9 | No growth | Nil |
